# A Novel Aptamer-Based Approach for Lipoprotein Removal to Achieve Ultra-Pure Blood EV Isolation

**DOI:** 10.1101/2025.03.29.646082

**Authors:** Soyoung Jeon, YongWoo Kim, Sehyun Shin

## Abstract

Blood extracellular vesicles (EVs) are nanoscale lipid-bilayer particles that carry proteins, nucleic acids, and metabolites, rendering them powerful tools for non-invasive liquid biopsy and targeted drug delivery. However, clinical translation of blood-derived EVs is severely limited by the co-isolation of lipoproteins—whose size (30–100 nm) and density overlap with EVs—resulting in contaminated preparations that compromise biomarker accuracy and jeopardize therapeutic safety, efficacy, and biodistribution. To overcome this critical bottleneck, we developed ApoFilter, an aptamer-based affinity filtration platform engineered to selectively capture ApoB100- and ApoA1-containing lipoproteins (VLDL, LDL, HDL) while preserving EV integrity. In solutions containing only mixed lipoproteins, ApoFilter demonstrated a capture efficiency of 97% for ApoB100- and ApoA1-containing particles, yielding filtrates with >99% depletion of target lipoproteins. Crucially, this high selectivity was preserved in human plasma—a complex protein milieu—where ApoFilter removed over 99% of lipoproteins without any detectable loss of EV yield. Furthermore, when size-exclusion chromatography (SEC) or ExoTFF was integrated with ApoFilter, the limitations of each technique were innovatively complemented, achieving a highly advanced level of separation and purification. Notably, the ApoFilter–ExoTFF combination delivered the highest performance, attaining 99.9% protein removal and a 98.2% EV recovery rate, enabling ultrapure EV isolation from blood plasma. This synergistic purification strategy addresses both analytical and therapeutic requirements by eliminating lipoprotein interference in downstream molecular profiling and minimizing off-target effects in drug delivery. ApoFilter thus represents a versatile, scalable solution for isolating clinically relevant EVs from blood, substantially improving the reliability of liquid biopsy assays and accelerating the development of EV-based therapeutics.

## 1. Introduction

Extracellular vesicles (EVs) are nanoscale, lipid-bilayer particles (30–300 nm in diameter) secreted by virtually all cell types and enriched with proteins, lipids, mRNA, miRNA, and DNA that reflect the physiological state of their cells of origin (Hu et al. 2025). Encapsulation within a protective membrane preserves EV cargo integrity in complex biological fluids such as blood, enabling stable transport and uptake by recipient cells where they modulate signaling pathways and gene expression (Iannotta et al. 2021). These properties have propelled EVs to the forefront of precision medicine as both sensitive biomarkers and versatile therapeutic vehicles (Jia et al. 2022).

EV-based liquid biopsy has emerged as a minimally invasive approach for early disease detection and longitudinal monitoring across a spectrum of pathologies—including cancer, cardiovascular disease, and neurodegeneration—by profiling EV-associated nucleic acids and proteins in blood, urine, or saliva (Karimi et al. 2018). Compared with tissue biopsy, EV analysis offers repeatable sampling, reduced patient risk, and the potential to capture tumor heterogeneity in real time. Beyond diagnostics, EVs hold considerable promise as targeted drug delivery platforms due to their innate tropism for specific cell types and low immunogenicity, and they are under active investigation as modulators in immunotherapy (Khan et al. 2024).

Despite this potential, clinical implementation of EV technologies is fundamentally constrained by the co-isolation of blood lipoproteins—very-low-density (VLDL), low-density (LDL), and high-density (HDL)—which outnumber EVs by orders of magnitude and share overlapping size (30–100 nm) and buoyant density profiles (Kuma et al. 2024). Conventional isolation methods such as ultracentrifugation (UC), density gradient centrifugation (DGC) (Li et al. 2018), and size-exclusion chromatography (SEC) cannot fully resolve this overlap, resulting in preparations contaminated with lipoproteins that distort EV-specific protein and RNA signatures (Li et al. 2021). This contamination undermines the sensitivity and specificity of EV-based biomarkers, risking false positives or negatives that could misinform clinical decision-making. In therapeutic contexts, residual lipoproteins can alter EV biodistribution, reduce targeting efficiency, and provoke unintended immune responses, compromising both safety and efficacy(Monguió-Tortajada, Gálvez-Montón et al. 2019). Thus, the removal of lipoprotein contamination remains one of the most significant challenges in EV research and its clinical applications.

Immunoaffinity-based approaches utilizing aptamers—short oligonucleotides with high target specificity and chemical stability—have been extensively studied for the selective isolation of biomolecules from complex biological fluids (Phillips et al. 2021). In particular, aptamer sequences targeting ApoB100, a biomarker for (V)LDL, and ApoA1, a biomarker for HDL, have been identified, and their application in lipoprotein capture has been successfully demonstrated in previous studies (Sun et al. 2014).

In this study, we developed ApoFilter, a novel filtration system designed to address the challenge of lipoprotein contamination in plasma EV preparations by selectively capturing ApoB100- and ApoA1-containing lipoproteins. To achieve this, we functionalized a previously validated high-throughput, mesh-based filtration platform (Lee et al., 2025) with aptamers targeting ApoB100 and ApoA1, enabling highly efficient and selective lipoprotein depletion.

Performance analysis demonstrated that ApoFilter effectively captured lipoproteins within the mesh structure, resulting in their near-complete depletion (>99%) in the permeate. Furthermore, integrating ApoFilter into standard EV isolation workflows not only removed over 99% of lipoproteins but also maintained or significantly enhanced EV recovery, demonstrating a synergistic effect. Thus, ApoFilter simultaneously addresses one of the most persistent challenges in EV isolation—lipoprotein contamination—while improving EV yield, achieving both purification and recovery goals. This breakthrough establishes ApoFilter as a robust and scalable platform for ultrapure EV isolation, making it highly suitable for downstream molecular analysis and therapeutic applications.

## 2. Results and Discussion

To address the long-standing challenge of lipoprotein contamination in EV isolation, we developed an aptamer-based affinity filtration system, ApoFilter, designed to selectively remove (V)LDL and HDL from plasma samples prior to downstream EV purification. Conventional isolation methods such as UC and SEC rely on size and density differences, which are insufficient for resolving lipoproteins from small EVs due to their overlapping biophysical characteristics (30–100 nm diameter; 1.02–1.10 g/mL density). To overcome this limitation, we introduced a third axis of separation—molecular affinity—by engineering a filtration platform functionalized with aptamers targeting ApoB100 and ApoA1, surface markers of (V)LDL and HDL, respectively. By incorporating these affinity ligands into a stacked mesh configuration, ApoFilter enables selective retention of lipoproteins while allowing EVs to pass through (Fig. 1a).

**Figure 1.**
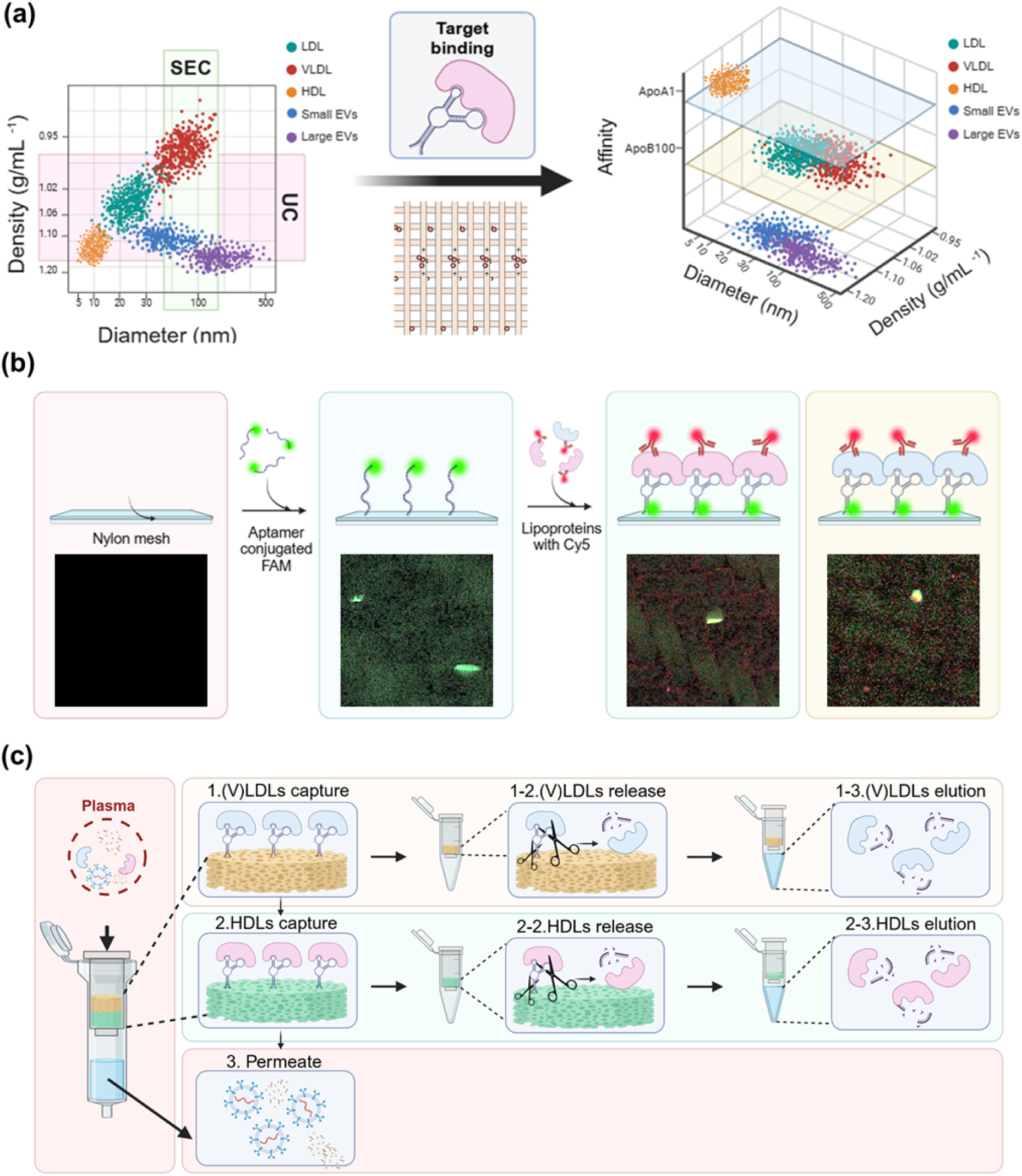
Aptamer-based affinity filtration enables rapid, selective removal of lipoproteins from plasma EV preparations. (a) Density vs. size distribution of plasma particles illustrates overlap between lipoproteins (LDL, VLDL, HDL) and EVs under conventional UC and SEC isolation; aptamer-functionalized mesh selectively captures lipoproteins via high-affinity binding; (b) Fluorescence microscopy confirms aptamer immobilization on nylon mesh (green) and specific binding of Cy5-labeled lipoproteins (red); (c) Workflow of sequential capture: (V)LDLs are first bound by ApoB100 aptamers, followed by HDL capture via ApoA1 aptamers, yielding a permeate enriched in EVs free of lipoprotein contamination.

To validate the functionalization and specificity of the system, we fluorescently labeled aptamers with FAM and confirmed their uniform immobilization on the nylon mesh via fluorescence microscopy (Fig. 1b). Subsequent incubation with Cy5-labeled lipoproteins resulted in co-localized green and red signals, confirming the successful and selective binding of (V)LDL and HDL to their respective aptamers.

The operational workflow involves sequential capture of (V)LDL and HDL using a two-layered mesh system. Plasma is first passed through an upper layer functionalized with ApoB100 aptamers, which captures (V)LDL. Bound particles are then enzymatically released via DNase I digestion and collected in an elution fraction. The filtrate continues to a second layer functionalized with ApoA1 aptamers for HDL capture, followed by a similar release and elution step. The final permeate contains EVs with lipoproteins effectively removed, setting the stage for high-purity downstream applications. The modular design of the filtration system enables effective demonstration of lipoprotein removal.

Figure 2 presents the validation of ApoFilter’s lipoprotein capture performance using purified (V)LDL and HDL solutions. To ensure precise assessment of target specificity, we employed defined lipoprotein standards instead of complex plasma samples. Each ApoFilter variant was applied to the corresponding lipoprotein type under previously described conditions as shown Fig. 2(a-c), and the capture efficiency was quantified using ELISA, nanoparticle tracking analysis (NTA), and BCA protein assays.

**Figure 2.**
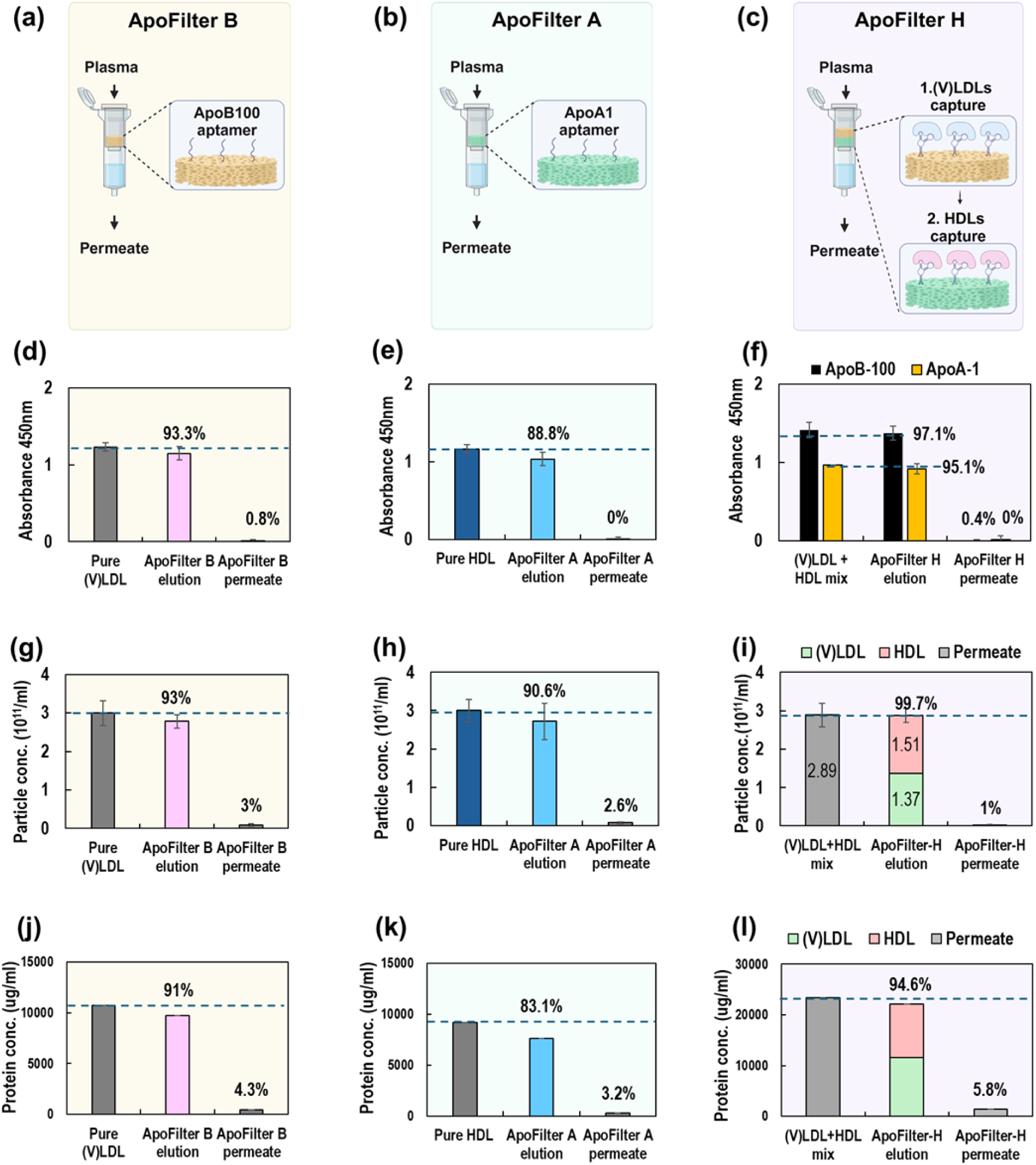
Validation of ApoFilter variants for selective lipoprotein capture using pure lipoprotein samples. (a–c) Schematics of ApoFilter B (ApoB100-aptamer), ApoFilter A (ApoA1-aptamer), and ApoFilter H (sequential (V)LDL then HDL capture), (d–f) ELISA quantification of lipoprotein levels in input, elution, and permeate fractions, demonstrating >88% capture efficiency and <1% residual lipoprotein in permeates, (g–i) Nanoparticle tracking analysis showing >90% depletion of lipoprotein particles from permeates, (j–l) Total protein measurements confirming >80% reduction in protein content in permeates for all filter types.

ELISA results (Figures 2d–f) demonstrated the high specificity of aptamer-mediated lipoprotein capture. For ApoFilter B, the absorbance at 450 nm decreased from 1.231 to 0.085, reflecting that just 0.8% of (V)LDL remained in the permeate. The capture efficiency in the elution buffer was 93.3%, showing a slight discrepancy compared to the minimal signal detected in the permeate. This may be attributed to reduced efficiency across the two-step process of target binding and enzymatic release. Simiarly, ApoFilter A, achieved 88.8% recovery, with no detectable HDL remaining in the permeate. When both lipoproteins were present (Figure 2f), ApoFilter H achieved 97.1% and 95.1% capture efficiency for ApoB100 and ApoA1, respectively, with less than 0.4% residual signal in the permeate.

Particle concentration analysis using NTA further confirmed these findings (Figs. 2g–i). ApoFilter B effectively removed approximately 97.0% of (V)LDL particles, as evidenced by the reduction of particle count in the permeate to just 3.0% of the original input (Fig. 2g). ApoFilter A achieved effective removal of HDL particles, with the particle count in the permeate reduced to 2.6% of the input, corresponding to a removal efficiency of approximately 97.4% (Fig. 2h). ApoFilter H achieved efficient removal of both lipoproteins from the mixed sample, leaving only 1.0% of particles in the permeate, which corresponds to a 99.0% overall removal efficiency (Fig. 2i). BCA protein quantification showed similar patterns to the NTA results, further supporting the effective removal of lipoproteins by ApoFilter.

In this experiment, human plasma samples were tested using three ApoFilter types. ELISA analysis of elution fractions (Fig. 3d) demonstrated high capture specificity. ApoA1, the HDL marker, was highly enriched in elution samples from both ApoFilter A and ApoFilter H, while ApoB100, the (V)LDL marker, was prominently detected in elution fractions from ApoFilter B and ApoFilter H. These results confirm that aptamer-functionalized meshes retain their selective binding capacity even in plasma, effectively isolating their respective lipoprotein targets. In contrast, EV markers such as CD9, CD63, and CD81 showed negligible ELISA signals in elution samples, indicating that EVs were not retained by the filters. In the permeate fractions (Fig.3e), ELISA analysis confirmed effective lipoprotein depletion, with ApoA1 nearly absent in ApoFilter A and H, and ApoB100 nearly absent in ApoFilter B and H, demonstrating the high specificity and efficiency of the ApoFilters.

**Figure 3.**
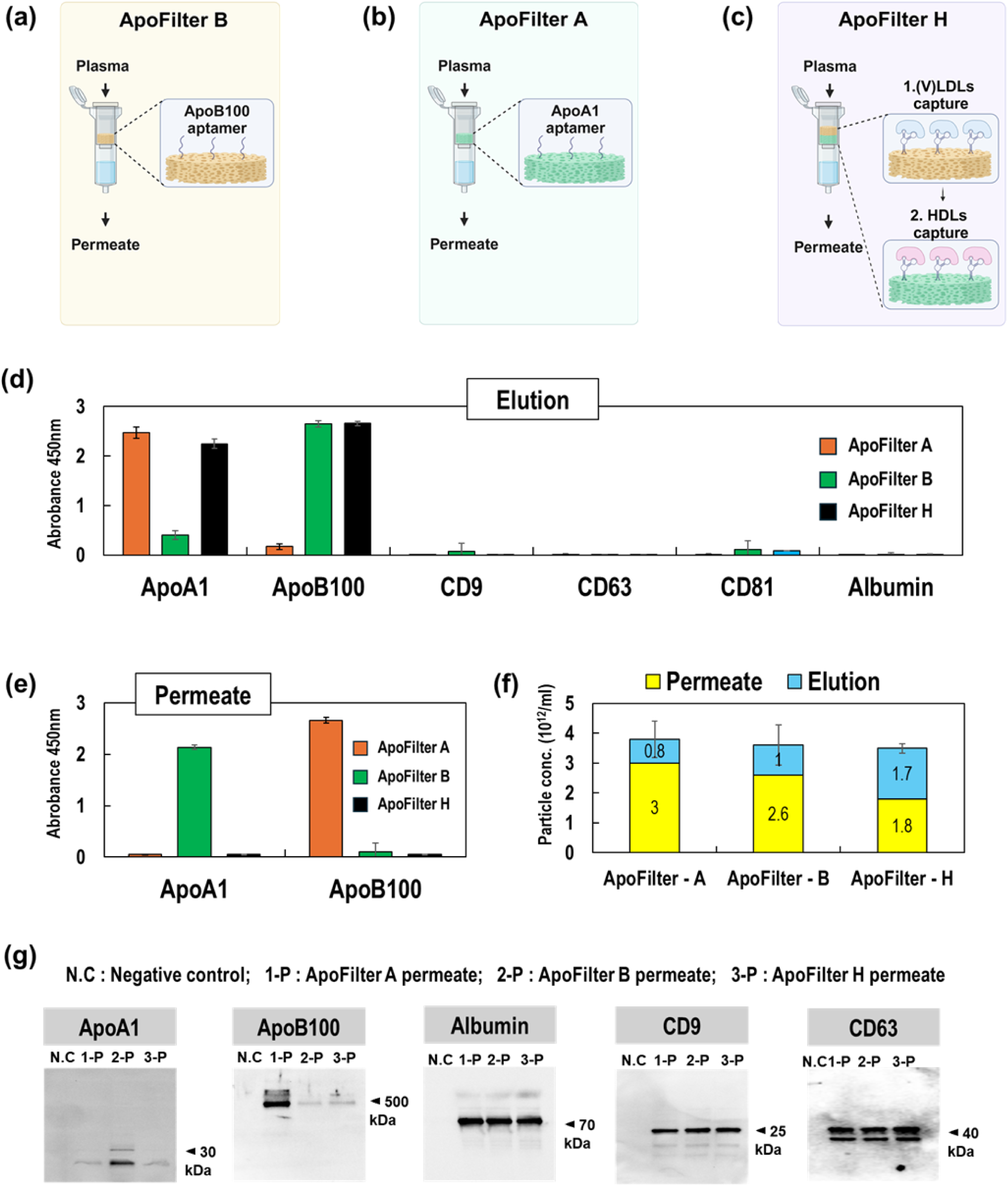
Performance Analysis of ApoFilter H for Blood Plasma. (a) Test Protocols, (b) Western blot analysis of permeate samples from different ApoFilter types, (e) ELISA analysis of permeate samples from different ApoFilter types, (f) NTA analysis of permeate/elution samples from different ApoFilter types, (g) ELISA analysis of elution samples from different ApoFilter type, (h) BCA assay of permeate/elution samples from different ApoFilter types.

The particle concentrations measured in the elution fractions of each filter were 0.8 × 10¹¹, 1.0 × 10¹¹, and 1.7 × 10¹¹ particles/mL for ApoFilter A, B, and H, respectively, with the combined total of A and B closely matching the value obtained from ApoFilter H. This indicates that ApoFilter H effectively captures both HDL and (V)LDL simultaneously. A similar trend was observed in the permeate fractions. Notably, the particle concentration in the permeate from ApoFilter H accounted for only 51.4% of the total particles. This suggests that approximately half of the particles detected by NTA in unfiltered plasma samples originate from lipoproteins rather than EVs.

Western blot analysis of permeate fractions (Figure 3g) supported the ELISA findings. ApoA1 bands were absent in permeates from ApoFilter A and H, while ApoB100 bands were undetectable in permeates from ApoFilter B and H, confirming target-specific removal. In contrast, albumin and EV markers (CD9, CD63) remained detectable across all permeate samples (except the negative control), indicating that ApoFilter does not retain non-target plasma proteins or EVs.

These results confirm that the present ApoFilters enable highly selective and efficient removal of lipoproteins even from human plasma, which is highly complex biological fluids. Each filter specifically capture (V)LDL and HDL, respectively under various proteins and their sequential integration in ApoFilter-H allowed for the simultaneous depletion of both lipoprotein subtypes within a single, modular device. Importantly, the filters maintained high performance in, successfully discriminating lipoproteins from non-target components such as EVs and plasma proteins.

Until now, we have evaluated the standalone performance of ApoFilter in removing lipoproteins, either from pure lipoprotein solutions or directly from plasma. In the following section, we assess the performance of ApoFilter when integrated with conventional EV isolation methods, evaluating not only its lipoprotein depletion capability but also its impact on EV recovery efficiency. For this investigation, we selected three commonly used EV isolation techniques: ultracentrifugation (UC), size exclusion chromatography (SEC), and ExoTFF. ExoTFF, recently developed by our group (Kim et al, 2025), combines electrokinetic separation principles derived from our previously reported ExoFilter technology with tangential flow filtration (TFF), which allows particles and fluid below a certain size threshold to pass through. Plasma samples were processed under six different conditions: each EV isolation method alone and each preceded by ApoFilter-H pretreatment, in order to compare the relative performance of each approach.

First, Western blot analysis was performed to detect major plasma protein markers in permeate samples obtained from each method (Figure 4(a)). Clear differences were observed in the ApoB100 and ApoA1 bands depending on whether ApoFilter H was applied. In contrast, other plasma proteins such as albumin and extracellular vesicle (EV) markers, including CD9, CD63, and albumin, exhibited similar expression patterns regardless of ApoFilter filtration. These findings indicate that ApoFilter selectively removes lipoproteins from plasma without affecting other plasma proteins or EVs.

**Fig. 4.**
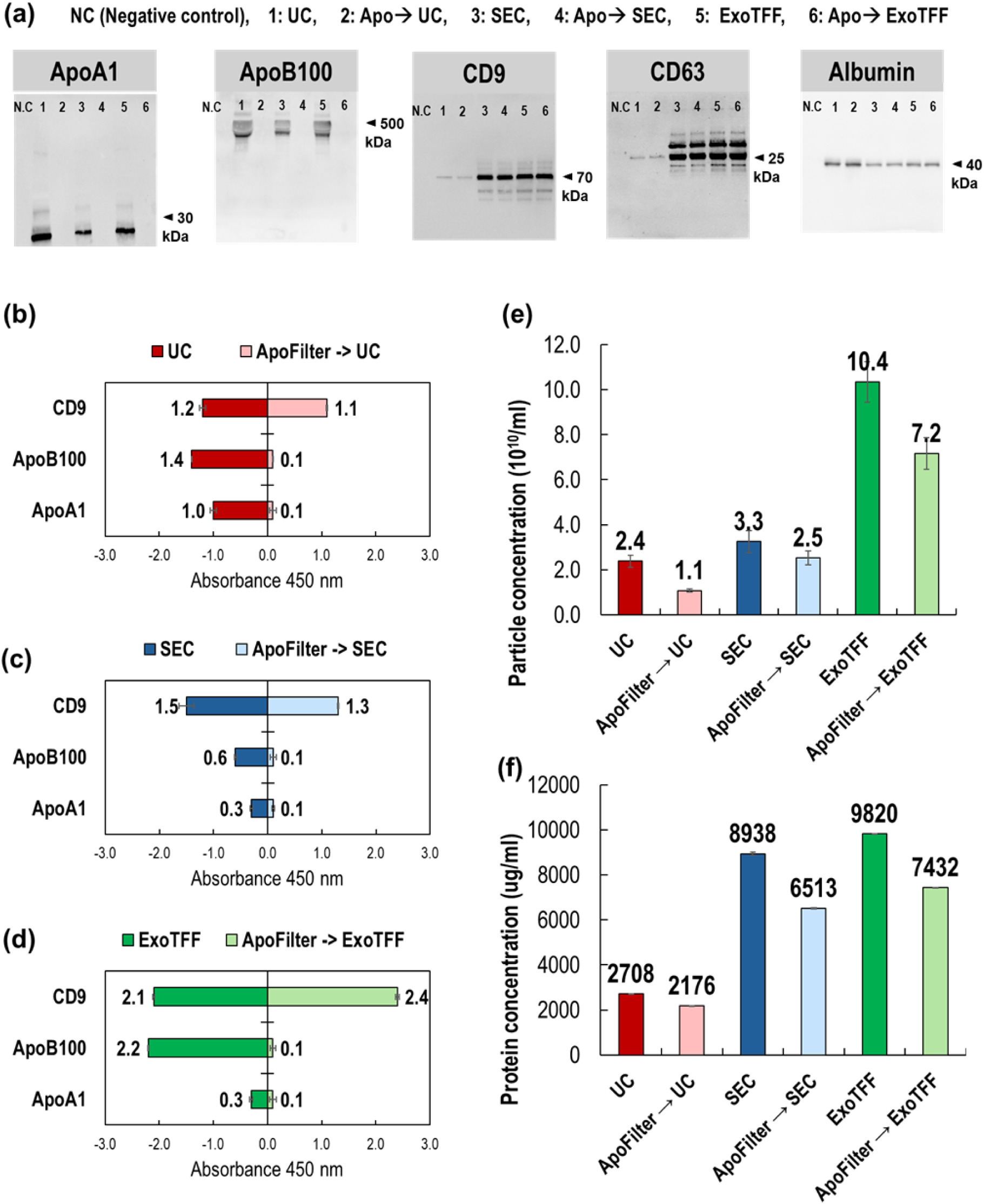
Synergistic removal of lipoprotein contaminants and preservation of EV yield by integrating ApoFilter with standard isolation workflows. (a) Western blot analysis of lipoprotein markers (ApoA1, ApoB100), EV markers (CD9, CD63), and albumin in samples isolated by ultracentrifugation (UC), size-exclusion chromatography (SEC), or ExoTFF with (+) or without (–) ApoFilter pretreatment, (b–d) ELISA quantification of ApoA1, ApoB100, and CD9 across each isolation workflow, (e) Nanoparticle tracking analysis showing particle concentrations for each method, (f) BCA assay of total protein concentrations for each workflow.

Next, ELISA results comparing lipoprotein and EV marker levels across different isolation methods are presented in Figure 4(b–d). When ultracentrifugation (UC) was applied alone (Fig. 4b), substantial levels of ApoB100 (1.4) and ApoA1 (1.0) remained, indicating incomplete removal of (V)LDL and HDL. However, with ApoFilter pretreatment (ApoFilter → UC), both markers were dramatically reduced to 0.1, demonstrating near-complete depletion. Meanwhile, CD9 absorbance showed minimal change, decreasing slightly from 1.2 to 1.1, suggesting that EV recovery was well preserved (91.7% retention).

In the case of size exclusion chromatography (SEC) (Fig. 4c), absorbance for ApoB100 and ApoA1 dropped to 0.6 and 0.3, respectively, indicating partial lipoprotein removal. Yet, when combined with ApoFilter, both markers were reduced to 0.1, again demonstrating the filter’s high depletion efficiency. CD9 absorbance changed modestly from 1.5 (SEC alone) to 1.3 (ApoFilter → SEC), reflecting a small reduction (about 13.3%) in EV recovery. Lastly, ExoTFF alone (Fig. 4d) resulted in higher CD9 absorbance (2.1), showing strong EV recovery but poor lipoprotein removal, with ApoB100 remaining at 2.2. When ApoFilter was applied beforehand (ApoFilter → ExoTFF), lipoprotein markers dropped sharply to 0.1, and CD9 increased to 2.4, indicating not only improved sample purity but also enhanced EV yield. These results confirm that ApoFilter selectively removes lipoproteins without compromising, and in some cases even enhancing, EV recovery—particularly when paired with ExoTFF.

When ExoTFF alone was employed, the reduction in ApoA1 signal was comparable to that observed with SEC, indicating a similar level of HDL removal. In comparison, the ApoB100 signal exhibited only a modest reduction, consistent with prior reports (Fisher et al. 2014), which indicate that despite a nominal TFF cutoff of 50 nm (800 kDa), larger lipoproteins like (V)LDL may not be completely removed. In contrast, both ApoA1 and ApoB100 markers were completely undetectable in the ApoFilter combined with ExoTFF sample. Interestingly, the CD9 absorbance was slightly higher in the ApoFilter + ExoTFF sample compared to ExoTFF alone. This suggests improved efficiency in targeting EV markers after removing lipoproteins and other impurities through ApoFilter.

Figure 4(e) presents the NTA analysis of particle concentrations across different EV isolation methods, with and without prior ApoFilter treatment. In all cases, samples that underwent ApoFilter pretreatment showed a marked reduction in particle concentration. For instance, particle counts decreased by approximately 54% for UC (from 2.4 × 10¹⁰ to 1.1 × 10¹⁰ particles/mL), 24% for SEC (from 3.3 × 10¹⁰ to 2.5 × 10¹⁰ particles/mL), and 30.8% for ExoTFF (from 10.4 × 10¹⁰ to 7.2 × 10¹⁰ particles/mL). These reductions in particle concentration can be interpreted as the degree of benefit gained from ApoFilter treatment—specifically, the removal of lipoproteins that would otherwise remain undetected by the conventional isolation methods alone. From this perspective, UC demonstrated the greatest improvement in purity, followed by ExoTFF and SEC.

This consistent reduction reflects the effective elimination of lipoproteins by ApoFilter—particles that are similar in size and optical properties to EVs and therefore easily miscounted in NTA measurements. Without prior removal of these contaminants, NTA data can be misleading, as lipoproteins significantly contribute to overall particle counts. Much like the saying ‘not all that glitters is gold,’ not all particles detected by NTA are extracellular vesicles. By removing this major source of interference, ApoFilter improves the specificity and reliability of NTA-based EV quantification.

Figure 4(f) shows total protein concentrations measured by BCA assay, highlighting the effect of ApoFilter pretreatment across different EV isolation methods. In all cases, samples processed with ApoFilter exhibited reduced protein levels compared to their non-filtered counterparts, reflecting the removal of lipoprotein-associated proteins. This trend closely mirrors the particle count reductions observed in the NTA results, further supporting the conclusion that ApoFilter effectively eliminates non-EV contaminants and improves overall sample purity.

Figure 5 provides a comparative overview of three commonly used EV isolation methods—ultracentrifugation (UC), size-exclusion chromatography (SEC), and ExoTFF—applied alone or in combination with ApoFilter, using plasma samples. Although plasma values are not visually presented in Figure 4(b–d), the original absorbance values of plasma samples were measured in parallel and served as reference points for calculating the removal efficiency of lipoproteins (ApoB100, ApoA1) and the recovery efficiency of EV markers (CD9). These calculated values allow a more accurate comparison of each isolation strategy’s performance and are visualized in the radar plots of Figure 5. EV recovery was assessed using three canonical EV markers (CD9, CD63, and CD81), while purification performance was evaluated based on the depletion of major plasma contaminants, including lipoproteins (ApoA1, ApoB100) and albumin. The radar plots clearly illustrate the trade-offs and synergistic effects across the evaluated workflows.

**Figure 5.**
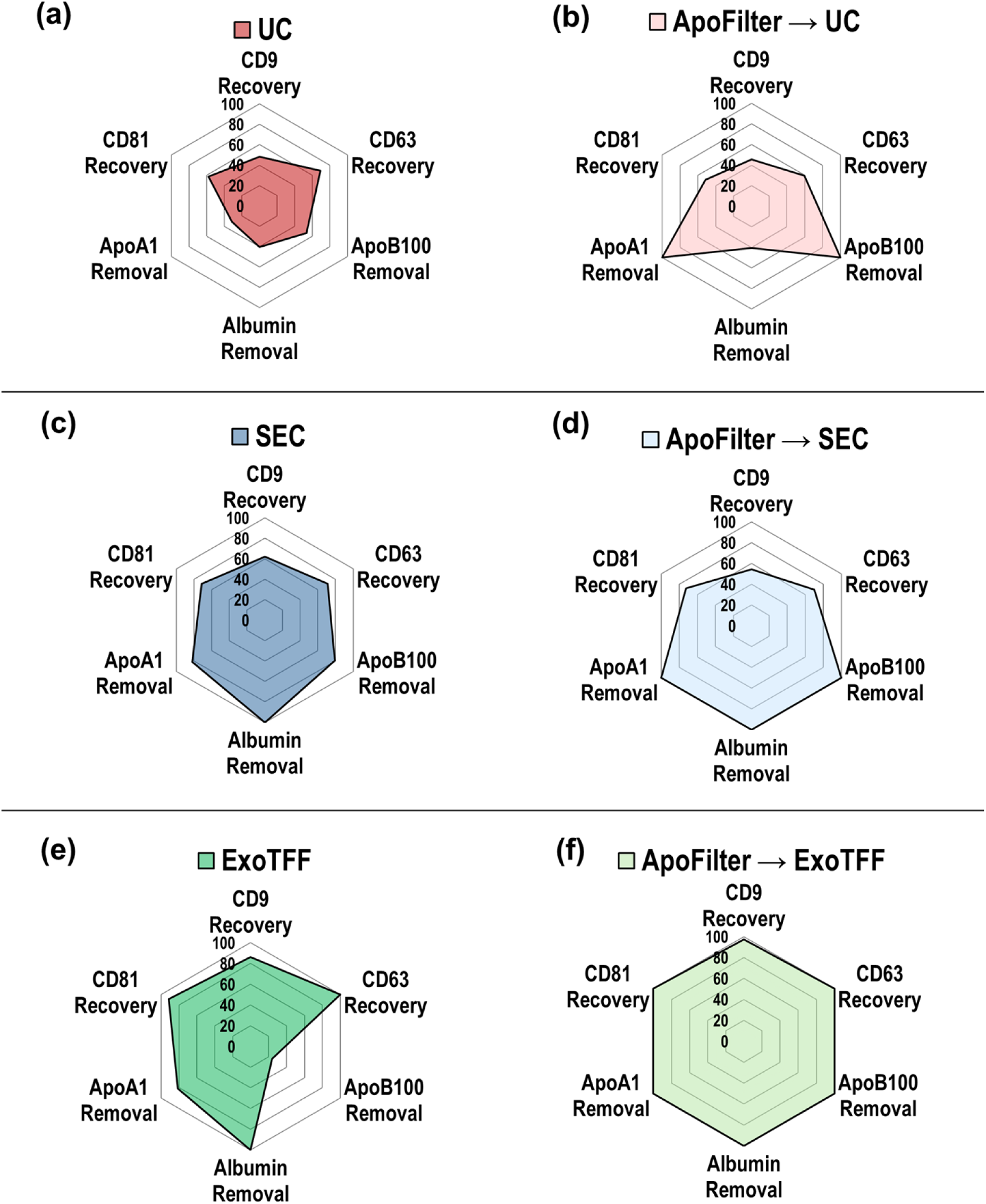
Performance comparison of EV isolation workflows with and without ApoFilter pretreatment. Radar plots depict percentage recovery of EV markers (CD9, CD63, CD81) and removal of contaminant proteins (ApoA1, ApoB100, albumin) for each method: (a) UC, (b) ApoFilter→UC, (c) SEC, (d) ApoFilter→SEC, (e) ExoTFF, and (f) ApoFilter→ExoTFF. Integration of ApoFilter markedly increases lipoprotein (ApoA1/ApoB100) and albumin depletion while maintaining or enhancing EV marker recovery across all workflows.

UC alone demonstrated modest EV recovery but poor impurity removal, with particularly low depletion rates for lipoproteins and albumin. Upon integration with ApoFilter, however, lipoprotein removal markedly improved, while EV recovery remained largely unaffected. SEC, in contrast, achieved relatively better performance in albumin removal and moderate lipoprotein depletion, albeit with slightly lower EV recovery. When combined with ApoFilter, SEC exhibited near-complete removal of both lipoproteins and albumin, with only a marginal impact on EV yield.

ExoTFF, known for its high EV recovery, showed limited capacity to remove larger lipoproteins such as (V)LDL. Remarkably, the combination of ApoFilter with ExoTFF not only enabled efficient removal of both ApoB100- and ApoA1-containing lipoproteins, but also enhanced EV recovery, highlighting a synergistic benefit. Collectively, these findings underscore the value of integrating ApoFilter into EV isolation workflows to simultaneously enhance purity and yield, especially in plasma samples where lipoprotein contamination poses a significant analytical challenge.

In this study, we developed and evaluated the efficacy of an aptamer-based affinity filtration technology (ApoFilter) designed to address the persistent issue of lipoprotein contamination in extracellular vesicle (EV) isolation from blood. Through extensive experimentation using mixed lipoprotein samples and plasma samples, we validated the high specificity and efficiency of ApoFilter. Particularly notable was its ability to achieve exceptionally high lipoprotein removal efficiencies, over 99%, resulting in significantly improved purity of isolated EV samples. hese findings are supported by Figure 5, which illustrates the superior particle and protein purity achieved when ApoFilter is integrated with ExoTFF.

A key advantage of the ApoFilter system lies in its operational simplicity and speed. Unlike conventional immunoaffinity methods that require prolonged incubation times (> 1 hr), ApoFilter achieves rapid target capture during brief, gravity-driven filtration ( < 1 min, without compromising binding efficiency. This accelerated kinetics can be attributed to the synergistic design features. First, the ApoB100- and ApoA1-targeting aptamers employed here possess sub-nanomolar affinity, enabling near-instantaneous binding upon molecular contact (Delcanale et al., 2020). Second, the mesh-based format enforces convective flow through a densely functionalized surface, converting diffusion-limited interactions into collision-driven capture events and dramatically increasing the frequency of aptamer–lipoprotein encounters (Lee et al, 2025). Finally, the multilayer architecture of the mesh exponentially expands the available binding surface area and residence opportunities, ensuring that even transient passage of lipoprotein particles results in high-efficiency capture (Lee et al, 2025). Collectively, these elements transform what is ordinarily a slow, diffusion-limited immunoaffinity process into a rapid, high-throughput filtration platform—resolving a key bottleneck in EV purification by coupling exceptional specificity with near-instantaneous performance.

ApoFilter is readily compatible with all conventional EV isolation workflows, enabling synergistic improvements in both purity and yield. Although our protocol applied ApoFilter as a pre-filtration step before ultracentrifugation (UC), size-exclusion chromatography (SEC), or ExoTFF, the order of operations can be adjusted to suit specific workflows. Notably, the greatest benefit was observed when ApoFilter was combined with ExoTFF: EV recovery increased from 85.4% to 98.2%. We attribute this enhancement to the immunoaffinity removal of lipoproteins—particles that share nearly identical size and surface charge with EVs—by ApoFilter, which produces a substantially cleaner feed for the downstream electrokinetic capture employed by ExoTFF (Kim et al, 2025). In contrast, density-based UC and size-based SEC showed minimal improvement in EV recovery even after ApoFilter prefiltration, indicating that lipoprotein contamination has little impact on these methods. Conversely, ExoTFF—which relies on electrokinetic isolation—is markedly enhanced by the prior removal of similarly sized, negatively charged lipoproteins, resulting in significantly improved EV recovery and purity.

However, this study presents several limitations that must be addressed in future research. Firstly, although ApoFilter showed impressive performance in plasma, further validation in serum, urine, or saliva would broaden its clinical applicability. Secondly, our evaluation focused primarily on the purity and recovery rate of EVs; future studies should further explore the functional characteristics of isolated EVs, including their role in intercellular signaling and immune modulation. Additionally, this study utilized only ApoB100 and ApoA1 aptamers; future research could expand the aptamer library to target a wider range of lipoproteins and interfering substances, potentially integrating them for improved efficiency. Lastly, the practical implementation of ApoFilter in clinical settings requires development of standardized protocols and an automated filtration system for high-throughput and consistent processing.

## 3. Conclusions

In conclusion, this study demonstrated the efficacy and utility of ApoFilter, an aptamer-based affinity filtration technology, as an innovative solution to the longstanding challenge of lipoprotein contamination in EV isolation from blood. ApoFilter H significantly enhances EV purity by selectively removing lipoproteins without affecting other critical plasma components, including EVs and associated biomarkers. This advancement promises to greatly enhance EV-based liquid biopsy applications, supporting early disease diagnosis, personalized medicine, and therapeutic monitoring. Future research addressing the identified limitations and optimizing ApoFilter H could further reinforce its potential as a robust, clinically applicable technology in EV-based diagnostics and therapeutics.

## 4. Materials and Method

### 4.1 Design of the Aptamer

The ApoB-100 aptamer (5’-ACCT CGAT TTTA TATT ATTT CGCT TACC AACA ACTG CAGA-C6-NH₂-3’), ApoA-1 aptamer (5’-CCTC GGCA CGTT CTCA GTAG CGCT CGCT GGTC ATCC CACA-C6-NH₂-3’) and ApoE aptmaer (5’-ACT AGC TAC GGG GTG GGT GGG CGG TGT CAG TTT GTT TAT TGG TGC TAT ACA TCC TCT ATA-C6-NH₂-3’) were synthesized by Bioneer Co. (Daejeon, Korea).

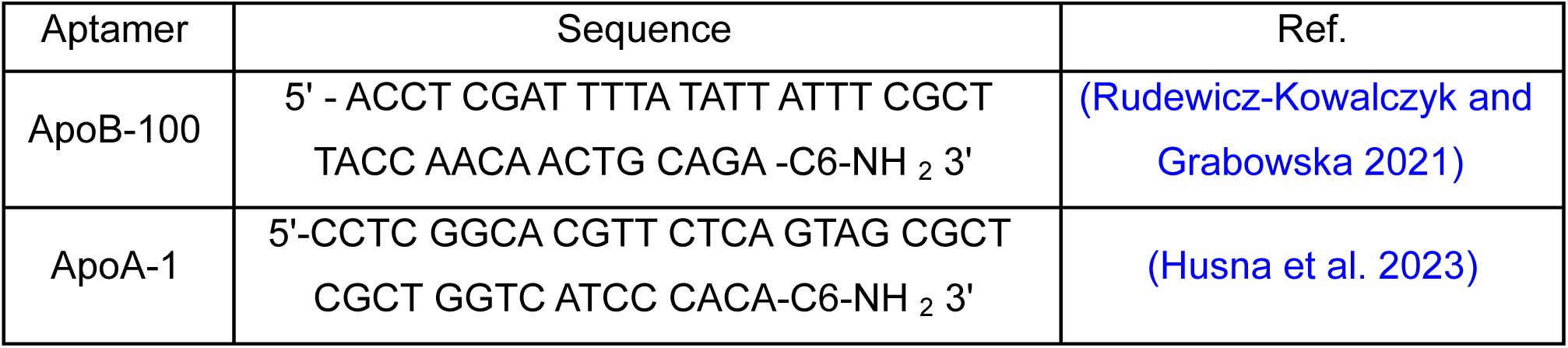

### 4.2 Preparation of aptamer coated mesh

Nylon meshes (Lixin Huarun MESH Co., China) were cut to a diameter of 11 mm using a laser cutter (BEAMO, MIRTECH Korea) and subsequently functionalized through aptamer conjugation. The process began by placing the mesh pieces in a Petri dish and adding 5 mL of 0.1N hydrochloric acid, followed by incubation for 30 minutes. After incubation, the meshes were washed three times with deionized (DI) water to remove residual acid. Subsequently, 10 mL of 2.5% glutaraldehyde solution (Sigma-Aldrich, USA) was added to the Petri dish, and the meshes were incubated for 30 minutes. After the incubation, the meshes were thoroughly washed with DI water to remove any residual glutaraldehyde. For the aptamer conjugation, 50 mg of nylon mesh was mixed with 1.4 μg of aptamer, 0.1M of 1-[3-(dimethylamino)propyl]-3-ethylcarbodiimide methiodide (EDC, Sigma-Aldrich, USA), and 0.1M of N-hydroxysuccinimide (NHS, Sigma-Aldrich, USA) diluted in 5 mL of 0.9% NaCl. The mixture was subjected to shaking on a roller mixer for 6 hours to facilitate the conjugation process. Finally, the meshes were washed three times with DI water to remove any unbound aptamers.

### 4.3 Preparation of various samples

This study adhered to the ethical standards of the Declaration of Helsinki. Plasma samples were sourced from Zen-Bio Inc. (Research Triangle, NC, USA). Additional samples, including cell culture media from umbilical cord mesenchymal stem cells, urine, and saliva, were collected in sterile containers and purified through centrifugation. Pure (V)LDL samples (#L8292, Merck KGaA, Darmstadt, Germany) and pure HDL samples (#L1567, Merck KGaA, Darmstadt, Germany) and plasma samples were centrifuged at 3000 × g for 15 minutes to sediment and remove large particulates. The clarified supernatant was then passed through an 800 nm pore-sized mesh to exclude particles larger than 800 nm. Samples were subsequently stored at -80°C pending further analysis.

### 4.4 Confocal imaging

Bare mesh, nylon mesh conjugated aptamer linked FAM (BioLegend, San Diego, CA), and nylon mesh captured lipoprotein linked Cy5 (BioLegend, San Diego, CA), with aptamer with FAM were imaged using a Zeiss LSM800, as indicated on individual figure legends. Images of lipoproteins were taken using a 100 × objective (LSM800; 100x = NA 1.4, LSM880; 100x = NA 1.46). Images of cells were captured using a 63 × objective (NA = 1.4). Images were acquired using 4 × line averaging. Z stacks were taken sectioning the height of the lipoproteins. Images were analysed using Zen black software (Zeiss) or ImageJ (FIJI). Colocalisation of dual labelled EVs was analysed through the JACOPx Plugin on ImageJ FIJI. Pearson’s coefficient, Rank-weighted colocalisation (RWC), fluorescent particle counts and colocalisation based on centres of mass-particles coincidence were performed using the JACOPx Plugin settings with manually adjusted thresholds, matching the intensity of particles seen on the software with the original image captured. The percentage of dual stained EVs was calculated using the object based ‘colocalization based on mass-particle coincidence’ feature of the JACOPx plugin. This automatically quantified the number of particles and assessed overlap of pixels calculated between the channels. The values of coincidence were then used to calculate the percentage of dual stained EVs as a proportion of the individual EV particles counted.

### 4.5 Scanning electron microscopy (SEM) Images

An anodic aluminum oxide (AAO) membrane mounted in a gasket was utilized for SEM measurements of EVs isolated from plasma. The samples were filtered through the membrane for SEM analysis. Following filtration, the membranes were incubated in a glutaraldehyde solution (Sigma-Aldrich, St. Louis, MO, USA) for 30 minutes. The membranes were then sequentially rinsed with 25%, 50%, 75%, 90%, and 100% ethanol and incubated in a dry oven at 37°C for 2 hours. After coating the membrane with platinum (Pt), the EVs and clusters present on the membranes were observed using an SEM (Quanta 250 FEG; FEI, Hillsboro, OR, USA).

### 4.6 Transmission electron microscopy (TEM) image

TEM was performed to visualize lipoproteins isolated using various methods, including the ApoFilter. A Caborn Formvar Film-150 copper grid was carefully handled with tweezers, ensuring the sample side was facing upwards, and placed on a Petri dish. Approximately 15 µL of the lipoprotein sample was applied to the grid and allowed to adsorb for 1 minute. To avoid contamination, the grid was covered during this incubation period. For negative staining, the grid was positioned vertically at a 90-degree angle, and 1% uranyl acetate was gently applied dropwise with a syringe, allowing the solution to flow downward. Excess staining solution was removed by blotting with filter paper. The grid was then left to air-dry on filter paper. After the drying process, lipoproteins were imaged using a JEM-1400 Flash (JEOL Ltd., Japan) transmission electron microscope. TEM images were captured at an accelerating voltage of 120 kV.

### 4.7 Nanoparticle Tracking Analysis (NTA)

Nanoparticle Tracking Analysis was performed utilizing the NS300 system along with NTA 3.4 Software (NanoSight, Wiltshire, UK) to evaluate both EVs and lipoproteins. The samples were first diluted in pre-filtered PBS. For analysis, three 30-second video recordings were made for each sample, with the camera level adjusted to 14 and the detection threshold set to 11. These recordings facilitated the capture and subsequent analysis of both EVs and lipoproteins, enabling the determination of their average size and concentration based on the dilution factors used.

### 4.8 BCA assay

Protein concentrations were determined using the Pierce™ BCA Protein Assay Kit (#23225; Thermo Scientific, Waltham, MA, USA). A calibration curve was prepared with bovine serum albumin, ranging from 0 to 2000 μg/mL, in nine incremental dilutions combined with the assay reagent. Triplicate measurements were conducted for each standard and sample. A volume of 100 μL from each sample was mixed with 2.0 mL of assay reagent and incubated at 37°C for 30 minutes. Post-incubation, samples were allowed to reach room temperature before measuring absorbance at 562 nm using a DS-11 spectrometer (Denovix, Wilmington, DE, USA). Protein concentrations were calculated from the standard curve by comparing the absorbance values of the samples to those of the blank standards.

### 4.9 Western blot assay

Protein extraction for Western blot analysis was performed on EVs and lipoproteins suspended in 200 µL of elution buffer. The extracted proteins were prepared by combining them with Laemmli buffer and 2-mercaptoethanol (BioRad, USA) and heated at 95°C for 10 minutes. Proteins were then isolated using SDS-PAGE on a Mini-PROTEAN® TGX™ Precast Gel (BioRad, USA). Western blotting was conducted using antibodies against specific EV markers CD9, CD81, TSG101, ALIX, and albumin as a negative control, as recommended by the MISEV 2018 guidelines (Théry et al., 2018). The antibodies used included recombinant anti-CD9 (ab92726, Abcam, Cambridge, UK), anti-CD81 (ab109201, Abcam, Cambridge, UK), anti-TSG101 (ab125011, Abcam, Cambridge, UK), and anti-ALIX (ab186429, Abcam, Cambridge, UK), along with goat anti-rabbit IgG H&L (ab205718, Abcam, Cambridge, UK). Visualization of protein bands was achieved using the ChemiDoc™ XRS+ System (Bio-Rad, CA, USA) with enhanced chemiluminescence (ECL) detection following the application of specific antibodies.

### 4.10 Immunocapture-based ELISA

Anti-CD9 (BioLegend, San Diego, CA), anti-CD81 (BioLegend, San Diego, CA), anti-ApoB100 (R&D Systems, Minneapolis, USA) and anti-ApoA1 (R&D Systems, Minneapolis, USA) antibodies were diluted to a concentration of 5 µg/mL in 10 mM PBS. 100µL of each diluted solution was added to microwells and incubated at 37°C for 2 hours. The wells were then washed with distilled water, followed by the addition of 200 µL of 0.5% casein in PBS, and incubated at 37°C for 1 hour. After additional washing, the 100µL samples were added to the wells, and the reaction was conducted 37°C for 2 hours. Following another rinse with distilled water, 100 µL of biotinylated anti-CD63 antibody (BioLegend, San Diego, CA),), biotinylated anti-ApoB100 (R&D Systems, Minneapolis, USA) and biotinylated anti-ApoA1 (R&D Systems, Minneapolis, USA) diluted to 1 µg/mL was added to each microwell for a 1hour reaction. After a subsequent wash, 100 µL of 0.45 µm membrane-filtered streptavidin-poly HRP20 (Fitzgerald, Acton, MA, USA) at 66 ng/mL was dispensed into each well and incubated at 37°C for 1 hour. To develop the signal, 200 µL of HRP substrate solution was added to the wells and incubated at room temperature for 15 minutes. The reaction was terminated by adding 50 µL of 2 M sulfuric acid to each well. The absorbance was measured at 450 nm using a microplate reader (SPECTROstar Nano, BMG LABTECH, Freiburg, Germany).

### 4.11 EV Isolation via UC

Ultracentrifugation (UC), though a demanding and time-intensive procedure requiring over 6 hours, is considered the gold standard for extracellular vesicle (EV) isolation. For this procedure, 1 mL of assorted biofluids was mixed with PBS at a ratio of 1:3 to facilitate EV isolation through UC. Initially, the samples underwent centrifugation at 3,000 × g for 15 minutes, followed by a subsequent spin at 12,000 × g for 30 minutes to remove larger vesicles and cell debris. The supernatant from this process was then ultracentrifuged at 120,000 × g at 4°C for 2 hours using a high-speed centrifuge (CP100WX; Hitachi, Tokyo, Japan). Post-ultracentrifugation, the supernatant was discarded, and the pellet was resuspended and further washed in PBS at 120,000 × g at 4°C for another hour. Finally, the pellet was resuspended in 200 µL of PBS, preparing it for further analysis.

### 4.12 EV Isolation via SEC

SEC columns used to isolate EVs was used the qEV columns (Izon Science). Blood collection and processing (≈20 min): blood is collected into plasma preparation tubes, mixed by inversion, and centrifuged 1,000 × g for 10 min at RT for plasma separation. Plasma preparation (≈15 min): plasma samples are filtered using 0.8-μm filters and centrifuged at 10,000 × g for 10 min at RT. The supernatant is collected to obtain filtered platelet-free plasma. EV isolation from plasma samples (≈25 min): a) 900 μL of filtered platelet-free plasma is loaded into the size exclusion chromatography column; b) F8(3.5–4.0 mL), F9(4.0–4.5 mL), and F10(4.5–5.0 mL) are collected; c) enriched EVs F8(3.5–4.0 mL), F9(4.0–4.5 mL), and F10(4.5–5.0 mL) are pooled in a final volume of 1,500 μL.

### 4.13 EV Isolation via ExoTFF

The ExoTFF system was applied to isolate EVs from various samples such as plasma, urine, saliva, and cell culture media (CCM). In the sample filtration step, the syringe-type ExoFilter is loaded with a 10 mL sample and then attached to the Tangential Flow Filtration (TFF) system. An empty syringe is attached to the opposite side of the TFF. The pistons of both syringes are used to perform repetitive movements. This configuration selectively captures negatively charged EVs within the positively charged mesh of the syringe-type ExoFilter, while impurities such as lipoproteins and proteins smaller than 30 nm are removed through the TFF along with the liquid components. In the elution step, which is the core process of ion exchange, 10 mL of 1M NaCl is used into the syringe-type ExoFilter to elute the EVs captured in the mesh, and the process is repeated until all the liquid is removed through the TFF. In the final recovery step, the syringe performing the role of the ExoFilter is replaced with a new syringe containing 2 mL of recovery buffer (PBS). This is cycled through the system to recover the EVs, resulting in a final product where the EVs are concentrated by approximately 1/5.

