## Supplementary information for "A Novel Aptamer-Based Approach for Lipoprotein Removal to Achieve Ultra-Pure Blood EV Isolation"

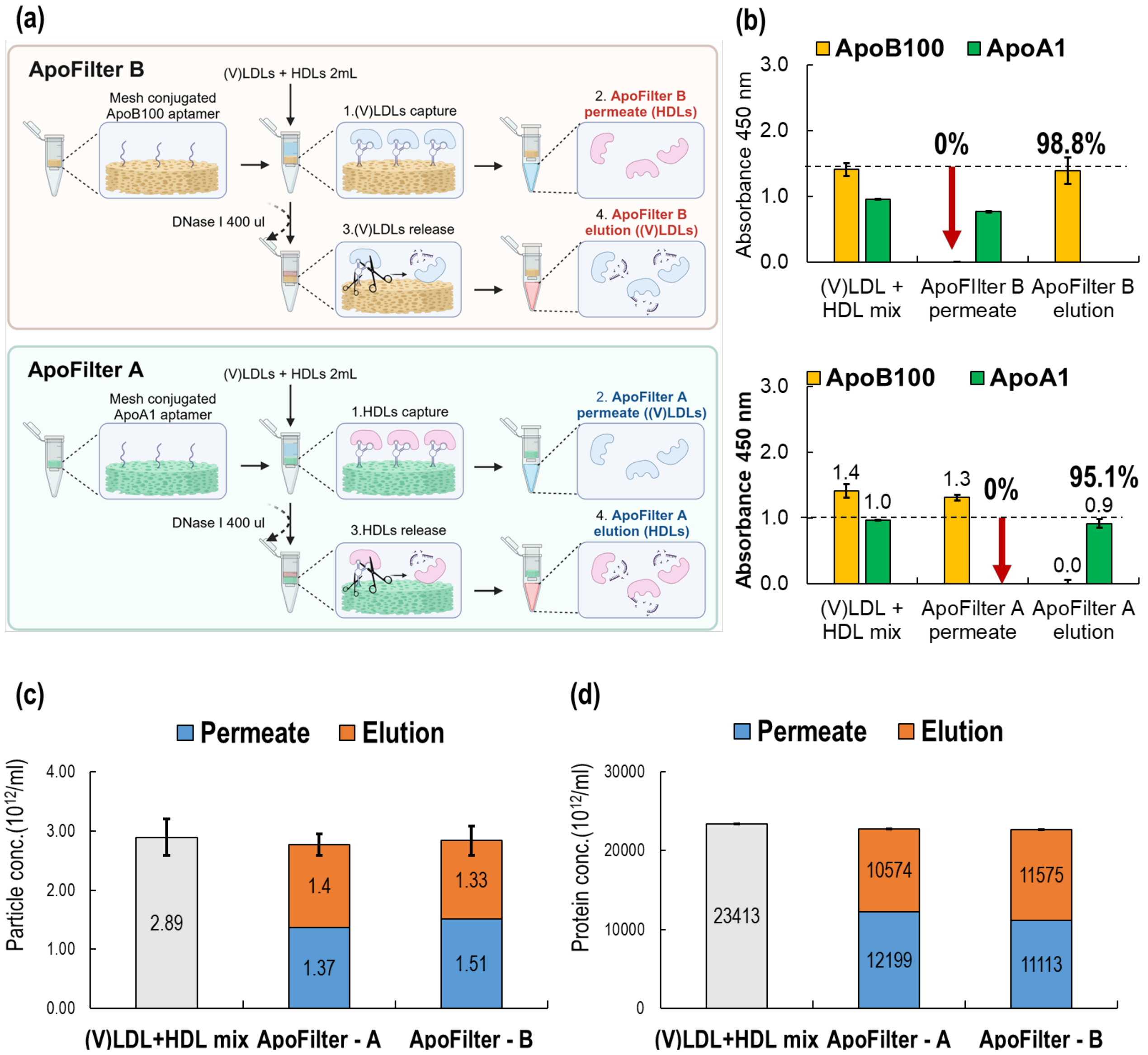


**Figure S1. Demonstration of ApoFilter's Selective Protein Capture ability in a Mixed Sample.** (a) Schematic Representation of ApoFilter B/A Permeate and Elution Sample Concepts, (b) TEM images of pure (V)LDL, pure HDL, (V)LDL + HDL mix, and ApoFilter B/A permeate/elution samples, (c) ELISA analysis of ApoFilter B/A permeate and elution samples, (d) NTA analysis of ApoFilter B/A permeate and elution samples, (e) BCA assay of ApoFilter B/A permeate and elution samples.

Figure S1 demonstrates the experimental results evaluating ApoFilter's ability to selectively capture target lipoproteins from a mixture of pure (V)LDL and HDL solutions. To assess this, we conducted experiments where mixed solutions of pure (V)LDL and HDL were separately applied to either ApoFilter B or ApoFilter A. For example, Figure S1(a) illustrates a case where a 2 mL mixed solution of pure (V)LDL and HDL was introduced into ApoFilter B. By design, (V)LDL particles were selectively captured by aptamers as the mixture passed through the filter (step 1), while the uncaptured HDL passed through the mesh and collected at the bottom of the column as the "permeate sample" (step 2). Subsequently, the captured (V)LDL particles were reversibly released using DNase I (step 3) and recovered via centrifugation (step 4). ApoFilter A operates similarly, selectively capturing HDL, while (V)LDL remains in the permeate.

ELISA assays were conducted to analyze the capture efficiency of ApoFilters (Figure S1(c)). For the original (V)LDL + HDL mixture, absorbance values were 1.411 for (V)LDL and 0.963 for HDL. In ApoFilter B permeate samples, (V)LDL was undetected (0%), whereas HDL detection was confirmed. Conversely, the elution samples from ApoFilter B showed a 98.8% recovery of (V)LDL with no HDL detected (0%). These results demonstrate that ApoFilter B, designed for selective capture of (V)LDL, successfully achieved its intended function in mixed samples.

Similarly, ApoFilter A, designed specifically for HDL capture, showed approximately 9% HDL detection in the permeate sample, whereas the elution sample showed a 95.1% recovery of HDL without any detectable (V)LDL. Thus, ApoFilter A demonstrated effective selective capture of HDL as originally designed.

Particle analysis using NTA further supported these findings (Figure S1(d)). The particle concentration of the (V)LDL + HDL mixture was $2.890 \times 10^{11}$ particles/mL. For ApoFilter A, particle concentrations in the permeate and elution samples were $1.370 \times 10^{11}$ (49.4%)

and 1.400 × $10^{11}$ particles/mL (50.6%), respectively, totaling 2.770 × $10^{11}$ particles/mL, corresponding to 95.8% recovery. Similarly, for ApoFilter B, the particle concentrations were 1.510 × $10^{11}$ (53.1%) in permeate and 1.330 × $10^{11}$ particles/mL (46.9%) in elution, totaling 2.840 × $10^{11}$ particles/mL, corresponding to a 98.3% recovery. Both results showed no statistically significant difference compared to the original mixed sample.

Protein quantification using BCA analysis (Figure S1(e)) yielded similar results. The protein concentration of the original mixed solution was 23,413 µg/mL. For ApoFilter A, protein concentrations in the permeate and elution samples were 12,199 µg/mL (53.6%) and 10,574 µg/mL (46.4%), totaling 22,773 µg/mL (97.3% recovery). For ApoFilter B, protein concentrations were 11,113 µg/mL in permeate and 11,575 µg/mL in elution, totaling 22,688 µg/mL (96.9% recovery). Again, no statistically significant differences from the original sample were observed.

Summarizing the results obtained so far, this study successfully demonstrated that the aptamer-based ApoFilter can selectively capture target lipoproteins from samples containing mixed lipoproteins. Additionally, the efficient capture of target molecules within the short duration of fluid passage through the mesh highlights the potential for high-throughput applications. Therefore, this study aims to utilize ApoFilter technology to remove lipoproteins, which pose significant challenges in extracellular vesicle (EV) isolation from actual blood samples, thereby enabling the extraction of EVs with higher purity.

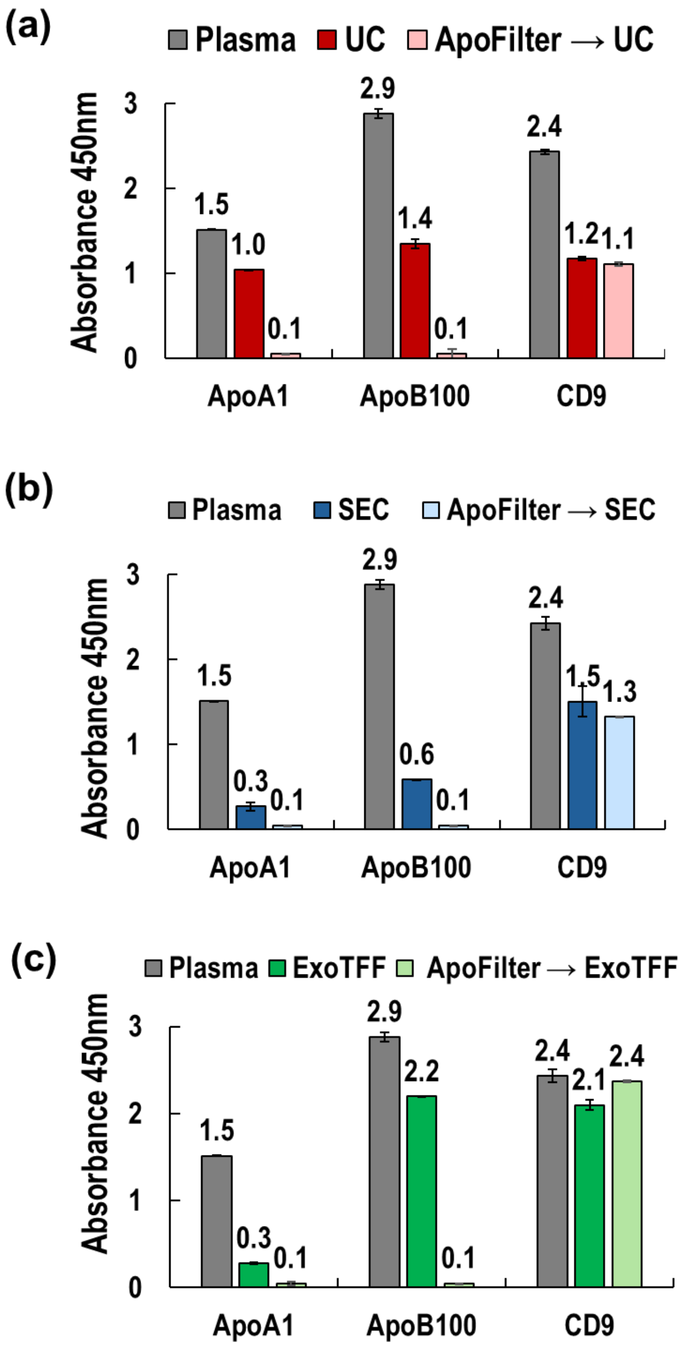


**Fig. S2 ELISA analysis of lipoprotein and EV marker levels before and after ApoFilter treatment across different EV isolation methods.** Absorbance at 450 nm was measured for plasma, conventional EV isolation methods (UC, SEC, ExoTFF), and their combinations with ApoFilter. (a) Ultracentrifugation (UC), (b) Size-exclusion chromatography (SEC), and (c) ExoTFF were evaluated with and without ApoFilter pre-filtration. Levels of ApoA1 and ApoB100 indicate the degree of lipoprotein removal, while CD9 reflects EV recovery. Plasma values were used as baseline references. ApoFilter pre-treatment significantly reduced lipoprotein markers across all methods without substantially compromising EV marker recovery.

Next, the ELISA results are presented in Figure S2. When ultracentrifugation (UC) was applied alone, the absorbance of ApoB100 decreased from 2.9 to 1.4 (48.3%), and that of ApoA1 decreased from 1.5 to 1.0 (66.7%), indicating a substantial amount of residual lipoproteins. This result further confirms the limitation of UC alone in effectively removing LDL and HDL. In contrast, when ApoFilter pretreatment was applied prior to UC, the lipoprotein markers were nearly undetectable. Additionally, the absorbance of the EV marker CD9 slightly decreased from 1.2 (UC alone) to 1.1 (ApoFilter + UC), corresponding to 91.7% retention. These results demonstrate that ApoFilter H enables near-complete lipoprotein removal—unachievable by UC alone—while maintaining a comparable level of EV recovery.

In the case of SEC (Fig.S2(b)), unlike UC, the absorbance levels of ApoA1 and ApoB100 significantly decreased compared to plasma, reaching 20.0% and 20.1%, respectively, but still exhibited residual lipoproteins. However, when ApoFilter was combined with SEC (ApoFilter + SEC), lipoproteins were no longer detectable. Consistent with previous results, CD9 absorbance was measured at 1.5 (62.5% relative to plasma) for the SEC-alone group and 1.3 (54.2% relative to plasma) for the ApoFilter + SEC group, indicating that the application of ApoFilter led to only an 8.3% reduction in EV recovery. This suggests that ApoFilter has minimal impact on EV yield.

As shown in Fig.S2(c), ExoTFF alone resulted in an ApoA1 reduction comparable to that of SEC, indicating similar HDL removal efficiency. In comparison, the ApoB100 signal exhibited only a modest reduction, which indicate that despite a nominal TFF cutoff of 50 nm (800 kDa), larger lipoproteins like (V)LDL may not be completely removed. In contrast, both ApoA1 and ApoB100 markers were completely undetectable in the ApoFilter combined with ExoTFF sample. Interestingly, the CD9 absorbance was slightly higher in the ApoFilter + ExoTFF sample compared to ExoTFF alone.

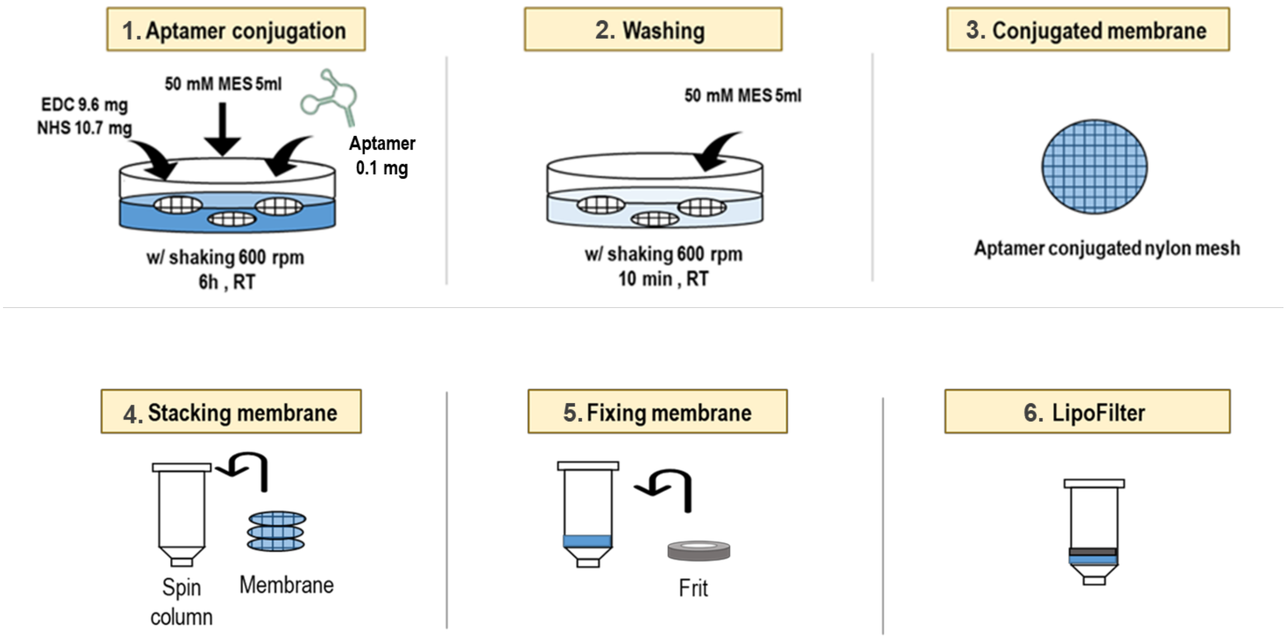


**Figure S3. Fabrication process of aptamer-conjugated nylon mesh for ApoFilter.** (1) Covalent conjugation of aptamers using EDC/NHS chemistry in MES buffer; (2) Post-conjugation washing with MES buffer to remove unbound reagents; (3) Resulting aptamer-conjugated nylon membrane; (4–5) Stacking of membranes into a spin column housing with frit support; (6) Assembly of the final LipoFilter device for lipoprotein capture.

To construct the ApoFilter, we developed a standardized protocol for conjugating aptamers onto nylon mesh membranes, enabling efficient and selective lipoprotein capture. As illustrated in Figure S3, aptamers targeting ApoB100 or ApoA1 were covalently immobilized onto the activated mesh surface using EDC/NHS chemistry in MES buffer. The optimal aptamer loading condition was determined to be 1.4 µg per membrane. After extensive washing to remove unreacted components, the aptamer-conjugated meshes were stacked into spin column housings with frit supports to form the final ApoFilter units. This modular assembly enabled consistent production of lipoprotein-selective filtration devices for use in downstream EV isolation workflows.
